# Modeling TCR–pMHC Binding with Dual Encoders and Cross-Attention Fusion

**DOI:** 10.64898/2025.12.01.691424

**Authors:** Wenbo Wang, Cong Qi, Zhi Wei

## Abstract

Accurately modeling the binding between T-cell receptors (TCRs) and peptide–MHC (pMHC) complexes is essential for guiding immunotherapy development and personalized vaccine design. However, the vast diversity of TCR repertoires and the scarcity of experimentally validated interactions make generalization to unseen epitopes challenging. This paper proposes TIDE, a cross-attention-driven dual-encoder framework that leverages large protein and molecular language models to learn discriminative representations of TCRs and peptides. In TIDE, TCR sequences are encoded using Evolutionary Scale Modeling (ESM), while peptides are transformed into SMILES strings and processed by MolFormer to capture chemical and structural properties. Multi-layer cross-attention then refines and integrates these embeddings, highlighting interaction-relevant patterns without requiring explicit structural alignment. Evaluated on the TCHard benchmark under both zero-shot and few-shot settings, TIDE achieves superior predictive accuracy and robustness compared to state-of-the-art baselines such as ChemBERTa, TITAN, and NetTCR. These results demonstrate that combining pretrained language models with cross-attention fusion offers a powerful approach for TCR–pMHC binding prediction and paves the way for more reliable computational immunology applications.

## I. Introduction

The recognition of antigens by T-cell receptors (TCRs) is a central process in the adaptive immune system, enabling the distinction between self and non-self and initiating targeted immune responses [1], [2]. Accurate modeling of TCR–peptide–MHC (pMHC) binding is critical for advancing immunotherapies such as PD-1/PD-L1 blockade, CAR-T therapy, and neoantigen-based vaccines [3], [4]. Predicting whether a TCR recognizes a specific peptide remains a significant challenge due to the vast diversity of TCR sequences and the structural complexity of peptide presentation by MHC molecules [5]–[10]. Traditional sequence-based methods often fail to capture subtle biochemical and spatial interactions, which limits their performance, particularly on rare TCRs or novel epitopes [11].

In computational studies, the complementarity-determining region 3 (CDR3) of the TCR *β* chain is the primary focus, as it plays a dominant role in peptide recognition and is consistently available in public datasets [12]–[16]. Following prior work [17], [18], this study leverages the CDR3 *β* sequence as the primary input for modeling TCR binding specificity. A key challenge in supervised modeling lies in generating negative samples for training and evaluation. Existing strategies include using curated non-binders from databases such as IEDB and NetTCR-2.0 [19] or constructing synthetic negatives through random or epitope-frequency-guided pairings [20], [21]. These approaches vary in biological plausibility and difficulty, making it essential to evaluate models across multiple negative sampling schemes to ensure robust benchmarking.

Recent advances in protein language modeling have transformed predictive modeling in immunoinformatics. Selfsupervised models such as TCR-BERT [22] and related pretraining approaches learn contextual and evolutionary features from large-scale TCR datasets, demonstrating strong generalization in low-data or zero-shot scenarios [23]–[25]. Building on these developments, the present work proposes a unified framework that integrates protein and molecular representations to model TCR–peptide interactions more effectively.

Accurate modeling of TCR–peptide interactions requires handling two distinct sequence modalities with very different length distributions. As shown in Figure 1, peptide sequences are typically short (7–16 amino acids, mean ≈ 10.7), whereas TCR *β* CDR3 sequences are longer and more variable (9– 23 amino acids, mean *≈* 14.5). This discrepancy poses a challenge for deep learning models, since short peptides may not provide sufficient sequential context to capture meaningful biochemical patterns.

**Fig. 1.**
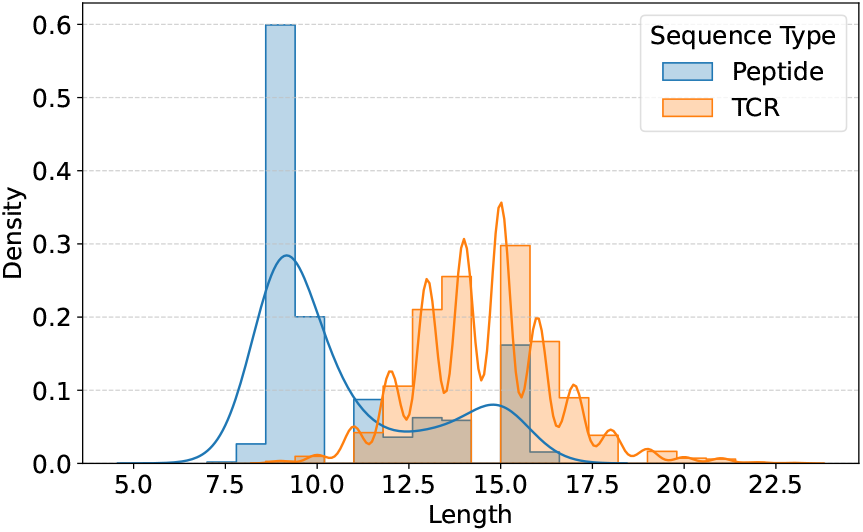
TCR and peptide sequence length distribution in the TCHard dataset.

This study introduces **TIDE** (TCR–pMHC Dual-Encoder), a two-stage deep learning framework designed to enhance the generalization of TCR–pMHC binding prediction. TIDE encodes TCR sequences using Evolutionary Scale Modeling (ESM) [26] to capture structural and evolutionary signals, while peptide sequences are represented in SMILES format and processed using MolFormer [28], which learns molecular graph features. This study makes the following key contributions:

### 1) TCR–pMHC binding prediction framework

We propose *TIDE*, a dual-encoder model leveraging ESM and MolFormer embeddings with Multi-Head Cross-Attention to capture TCR–peptide interactions.

### 2) Zero-shot and few-shot generalization

TIDE effectively generalizes to unseen epitopes with minimal supervision.

### 3) Comprehensive evaluation and analysis

We evaluate TIDE with multiple negative sampling strategies and embedding visualizations, showing that it yields biologically meaningful representations and outperforms baseline methods.

## II. Related Work

### Modeling TCR–pMHC Interactions

Computational prediction of TCR–peptide-MHC (pMHC) binding has evolved from similarity-based heuristics to deep learning models [8]. Early methods such as GLIPH [29] and TCRdist [30] cluster TCR sequences by CDR3 motif similarity, based on the assumption that structurally similar TCRs recognize the same epitopes. While these approaches are useful for discovering broad specificity groups, they rely on hand-crafted metrics and fail to capture complex nonlinear interactions. With the increasing availability of labeled data, machine learning methods emerged as a promising alternative. NetTCR [31] applies 1D convolutional neural networks (CNNs) to jointly encode TCR *β* CDR3 and peptide sequences, and NetTCR-2.0 [32] extends this approach with improved handling of variable sequence lengths. DeepTCR [16] further incorporates unsupervised representation learning to extract sequence motifs from both TCR and antigen repertoires. More recently, transformer-based architectures, such as TEINet [20], leverage pretrained encoders to capture long-range dependencies and improve generalization. These models have shifted the field toward end-to-end learning and demonstrated stronger performance on unseen epitopes, although most still rely on simple concatenation of TCR and peptide embeddings without explicit modeling of biochemical alignment.

### Sequence and Molecular Representations

Accurate representation of both TCR and peptide sequences is essential for reliable binding prediction. Most studies focus on the CDR3 region of the TCR *β* chain, which dominates antigen recognition and is well covered in public datasets [12]–[16], whereas the *α* chain is often omitted due to limited availability and smaller contribution [15]. Peptides are typically short (8–11 amino acids), which makes it challenging to capture their chemical and structural variability using raw sequences alone. To address this, recent work has explored molecular graph or SMILES-based representations [17], which retain information about bonding, functional groups, and spatial configuration. Meanwhile, self-supervised pretraining has emerged as an effective way to overcome data scarcity. Protein language models such as TCR-BERT [22], ESM [33], and ProtTrans [34] learn contextual embeddings from millions of sequences, improving generalization in low-resource settings [23]–[25]. Similarly, molecular models like MolBERT and MolFormer [28] leverage chemical structure pretraining to capture functional properties. These advances in sequence and molecular representation underpin modern TCR binding prediction frameworks.

## III. Methodology

The TIDE model adopts a two-stage deep learning framework that leverages transfer learning to capture the interaction patterns between T-cell receptor (TCR) sequences and epitope (peptide) sequences. Figure 2 provides an overview of the model architecture.

**Fig. 2.**
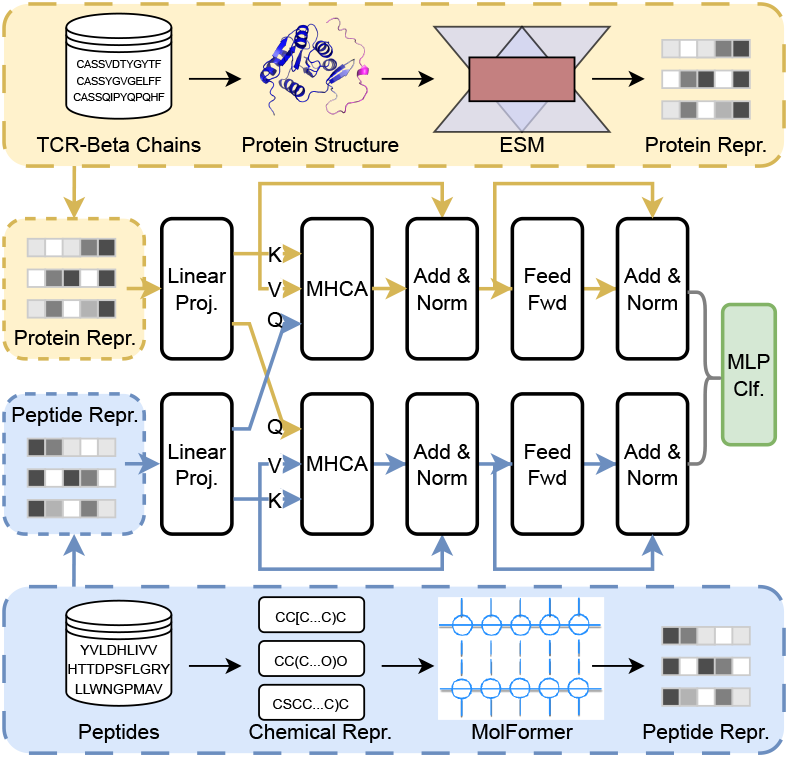
Framework of the TIDE Model. TIDE encodes TCR chains using ESM and peptide sequences via SMILES and MolFormer. A Multi-Head Cross-Attention module aligns TCR and peptide embeddings before classification. The model supports both random and reference-based negative sampling and generalizes effectively to unseen epitopes.

In the first stage, raw epitope sequences—typically short amino acid chains of around 10 residues—are converted into SMILES strings to mitigate the challenge posed by their limited length. This molecular representation produces sequences with an average length of approximately 185 characters, which is more suitable for transformer-based sequence modeling.

Subsequently, both TCR and epitope sequences are tokenized at the character level and processed by their respective pretrained encoders. These encoders project the sequences into low-dimensional vector embeddings that capture both structural and biochemical information. In the second stage, the resulting TCR and epitope embeddings are combined and passed through a fully connected neural network, which integrates information from both components to generate the final prediction.

### A. Sequence Encoding of TCRs and Peptides

The TCR (protein) and peptide (molecular) sequences are first encoded into vector embeddings using domain-specific pretrained encoders: ESM for TCR sequences and MolFormer for peptide SMILES strings.

#### TCR Encoding with ESM

Let the input TCR amino acid sequence be *S*_TCR_ = {*a*_1_, *a*_2_, …, *a*_*L*_}, where *L* is the sequence length. After residue-level tokenization, the sequence is embedded into a matrix *X* ∈ ℝ^*L×d*^, followed by Layer Normalization 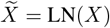 to stabilize training. Next, Multi-Head Self-Attention (MHSA) is used to model intra-sequence dependencies:

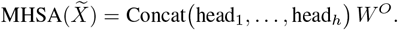

Each attention head is computed as

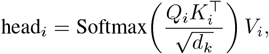

where the query, key, and value matrices are linear projections 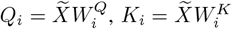 and 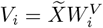.

Finally, a position-wise Feed-Forward Network (FFN) integrates local and global dependencies:

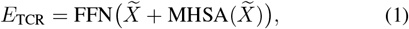

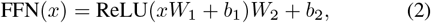

where *E*_TCR_ is the final TCR embedding.

#### Peptide Encoding with MolFormer

Given a peptide (epitope) sequence *S*_pep_, its molecular graph is first linearized into a SMILES string via depth-first traversal *S*_SMILES_ = DFS(*S*_pep_). The SMILES sequence is tokenized and embedded as a matrix *Y* ∈ ℝ^*T ×d*^, where *T* is its length. Layer Normalization and MHSA are then applied similarly to the TCR encoder:

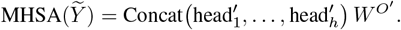

Each attention head is computed by

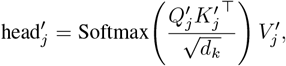

where 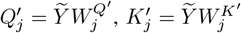, and 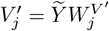.

The final peptide embedding is obtained using a feed-forward network:

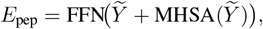

which integrates structural and chemical information for downstream cross-attention and classification.

### B. TCR–Peptide Interaction Modeling with MHCA

The Multi-Head Cross-Attention (MHCA) mechanism is designed to jointly capture the interactions between peptide 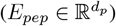 and TCR 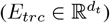 representations by leveraging bidirectional cross-attention followed by a compact MLP for final prediction. First, the peptide and TCR representations are projected to a shared latent space:

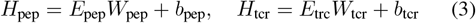

where 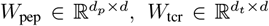 and *d* is the shared projection dimension.

To capture mutual dependencies between peptide and TCR embeddings, two cross-attention blocks are employed:

a. **Peptide attends to TCR:**

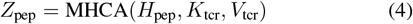

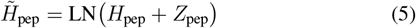

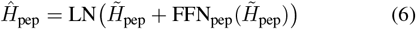
b. **TCR attends to Peptide:**

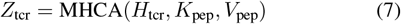

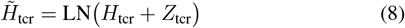

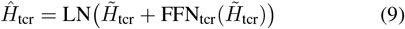

Here, FFN denotes a two-layer feed-forward network with GELU activations and dropout. Residual connections follow the standard Transformer architecture.

After removing the sequence dimension, the two attended embeddings are concatenated:

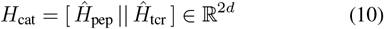

The concatenated representation is passed through a multilayer perceptron for final prediction:

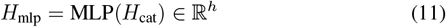

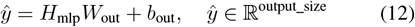

### C. Objective Function

The primary objective of the TIDE model is to predict the binary binding outcome between a TCR sequence and a peptide encoded as a SMILES molecule. The model outputs a scalar *ŷ* ∈ (0, 1), which represents the probability that a given TCR binds to a given peptide.

This problem is formulated as a binary classification task, and the Binary Cross-Entropy (BCE) loss is employed as the primary training objective:

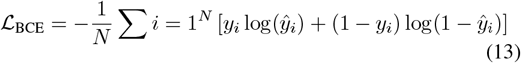

where *y*_*i*_ ∈ 0, 1 denotes the ground-truth label for the *i*-th TCR–peptide pair (1 for binding and 0 for non-binding), *ŷ*_*i*_ is the predicted binding probability from the final MLP layer, and *N* is the number of training examples.

An auxiliary regularization term is introduced to encourage alignment between *E*_trc_ and *E*_align_. This is implemented, for example, through a mean squared error constraint:

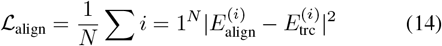

The overall training objective combines the classification loss and the alignment regularization term:

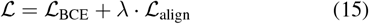

where *λ* is a tunable hyperparameter that controls the contribution of the alignment regularization term.

## IV. Experiments

To assess the performance of the TIDE model in TCR–peptide binding prediction, a set of experiments is conducted against state-of-the-art baselines across multiple datasets and evaluation settings. The experiments examine both zero-shot and few-shot learning scenarios, and further evaluate the quality of the learned sequence embeddings. The code and datasets for reproducing our experiments are available at: https://anonymous.4open.science/r/TIDE-4040

### A. Benchmark Evaluation of TIDE

The benchmark evaluation of TIDE aims to predict whether a given TCR binds to a specific peptide. In this framework, MolFormer is employed as the pretrained encoder for peptide SMILES sequences, while ESM is used for TCR sequences. The model is trained end-to-end using the Adam optimizer with early stopping based on the validation AUC. Depending on the experimental configuration, the pretrained encoders (ESM and MolFormer) are either kept frozen or fine-tuned, whereas the MHCA module and the MLP classifier are trained from scratch.

Model performance is rigorously evaluated on a held-out test set using metrics such as AUC-ROC to measure prediction accuracy and reliability. By combining the representational strengths of ESM and MolFormer, TIDE establishes a solid foundation for modeling protein–ligand interactions and provides insights that are valuable for drug discovery and biochemical research.

#### 1) Baselines

The baselines include **Large Language Model-based approaches**, including Smiles-Bert [35], TEINet [20], and ChemBERTa [36], as well as **end-to-end deep learning models** such as TITAN [17], ERGO II [7], NetTCR [32], Dlptcr [37], and Imrex [38].

#### 2) Dataset

The experiments utilize data derived from the TCHard dataset [19]. For all datasets, only the TCR *β* chain and the antigen peptide are used as model inputs, excluding TCR *α*, MHC, V, and J genes. This selection follows prior studies [39]–[41], which identify the *β* chain and the peptide as the most informative components for TCR–peptide interaction prediction.

### B. Negative Sampling Strategy

In TCR–peptide binding prediction, negative samples are essential for supervised training. Two common strategies are adopted for generating negative pairs: *reference control* and *random control* [19].

#### 1) Reference Control

This strategy constructs negative samples from experimentally validated datasets, where unbound TCR–peptide pairs are confirmed through biological assays. Although this approach provides highly reliable negative labels, it is resource-intensive and time-consuming, which limits its scalability to large datasets.

#### 2) Random Control

This strategy generates negative samples by randomly pairing TCR sequences with peptides. It is highly efficient and requires minimal additional resources; however, it introduces a risk of including pairs that may actually bind, potentially reducing the accuracy of the negative sample set.

#### 3) Negative Sampling Implementation

Among the negative sampling strategies, the random control is inherently more challenging, as it may inadvertently generate pairs that actually bind. To ensure that the model is evaluated on unseen peptides during testing (i.e., test peptides do not appear in the training set), the negative sampling procedure constructs a number of negative pairs equal to the positive pairs in the training data. Negative samples are generated by randomly pairing one peptide with one TCR from the training set, while preserving the original distribution of the TCHard dataset. This process produces the generated reference control dataset (Gen NA) and the generated random control dataset (Gen RN).

Formally, the negative sample set is defined as:

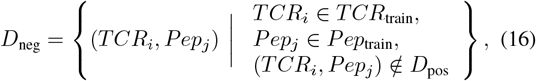

subject to

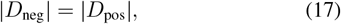

where *TCR*_train_ and *Pep*_train_ are the TCR and peptide sequences in the training set, *D*_pos_ is the set of positive training pairs, and *D*_neg_ is the generated negative sample set. Following this strategy, four datasets are used: the original reference control dataset (NA), the original random control dataset (RN), the generated reference control dataset (Gen NA), and the generated random control dataset (Gen RN).

All experiments employ a 5-fold split strategy. The original TCHard test set is used consistently across all folds, while the original training set is further divided into a new training set and a validation set in an 8:2 ratio. A strict *hard split* is maintained throughout, ensuring that no peptide in the validation or test set appears in the training set.

In total, the dataset contains approximately 160,000 unique TCR sequences and 1,000 unique peptides, resulting in roughly 220,000 TCR–peptide pairs in the training set, 60,000 in the validation set, and 40,000 in the test set.

### C. Performance Evaluation of TIDE

To rigorously evaluate the effectiveness of TIDE, we conduct a comprehensive comparison with state-of-the-art baselines across both the original datasets (Original NA and Original RN) and the generated datasets (Gen NA and Gen RN). The motivation for this experiment is twofold. First, benchmarking against diverse models, including both traditional deep learning and LLM-based methods, ensures a fair and transparent assessment of TIDE’s performance. Second, evaluating on both original and generated datasets allows us to examine how different negative sampling strategies impact the model’s ability to generalize to challenging TCR–peptide interactions.

Figure 3 presents the AUC-ROC scores of TIDE and baseline models across the NA and RN datasets. Overall, models exhibit noticeably stronger performance on the NA dataset, with AUC-ROC scores ranging approximately from 0.6 to 0.9, compared to 0.5 to 0.7 on the RN dataset. This trend indicates that the NA setting provides more favorable conditions for accurate TCR–peptide interaction prediction. When comparing model categories, LLM-based approaches consistently outperform conventional end-to-end deep learning models. The LLM-based models achieve higher AUC-ROC scores across both datasets, reflecting their stronger ability to capture the semantic and structural features of TCR–peptide interactions.

**Fig. 3.**
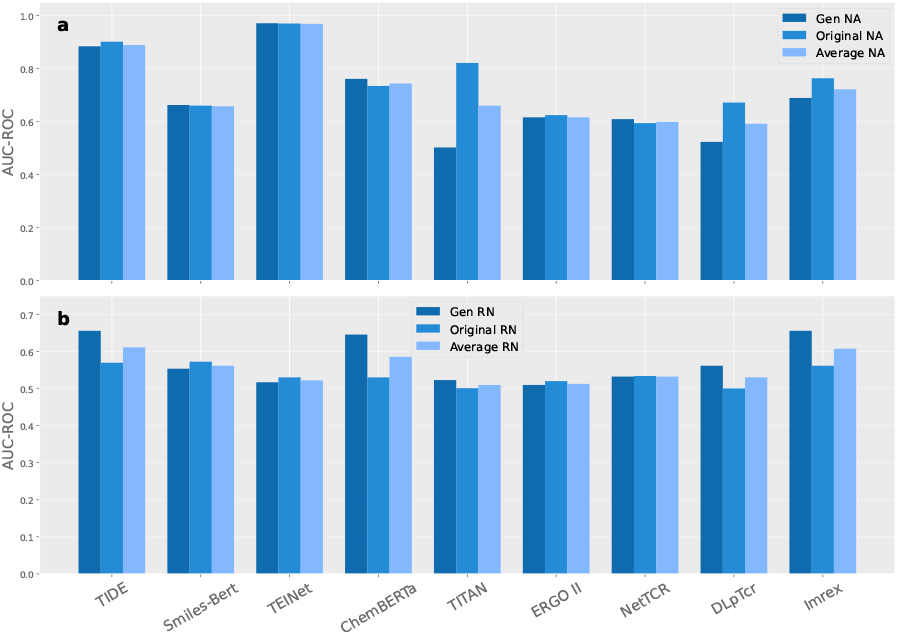
Performance of TIDE on different datasets. **a**. AUC-ROC scores across models for NA datasets. **b**. AUC-ROC scores across models for RN datasets.

TIDE demonstrates competitive or superior performance compared to most baselines. Although TEINet achieves slightly higher AUC-ROC on the NA datasets, our TIDE framework offers stronger interpretability and generalization. By integrating ESM and MolFormer embeddings with the MHCA module, TIDE explicitly models TCR–peptide interactions and captures structural and biochemical cues through SMILES-based peptide representations. These advantages enable TIDE to achieve high predictive accuracy while providing insights into the underlying binding mechanisms. This design enhances robustness in long-tailed or few-shot scenarios and offers biologically meaningful insights, as further demonstrated in the analyses presented in the following sections.

#### Impact of Negative Sample Generation on Model Performance

Table I reports the performance of all models across six evaluation metrics: AUC-ROC, Accuracy, Recall, Precision, F1-Score, and AUC-PR. Among these, AUC-ROC serves as the primary metric due to its robustness in binary classification settings.

**TABLE I.** Result of TIDE and baseline models on different datasets (Gen NA, Gen RN, Original NA, and Original RN).

| Model | AUC-ROC | ACC | RECALL | PRECISION | F1-SCORE | AUC-PR |
| --- | --- | --- | --- | --- | --- | --- |
| **Gen NA** |  |  |  |  |  |  |
| TIDE | 0.882 | 0.728 | 0.341 | 0.992 | 0.483 | 0.991 |
| Smiles-Bert | 0.659 | 0.702 | 0.621 | 0.843 | 0.811 | 0.808 |
| TEINet | 0.970 | 0.960 | 0.980 | 0.980 | 0.980 | 0.940 |
| ChemBERTa | 0.757 | 0.540 | 0.529 | 0.987 | 0.661 | 0.986 |
| TITAN | 0.502 | 0.620 | 0.523 | 0.590 | 0.608 | 0.970 |
| ERGO II | 0.613 | 0.665 | 0.618 | 0.873 | 0.702 | 0.698 |
| NetTCR | 0.605 | 0.683 | 0.592 | 0.793 | 0.721 | 0.688 |
| DlpTcr | 0.520 | 0.716 | 0.307 | 0.017 | 0.030 | 0.514 |
| Imrex | 0.685 | 0.448 | 0.433 | 0.985 | 0.593 | 0.986 |
| **Gen RN** |  |  |  |  |  |  |
| TIDE | 0.662 | 0.620 | 0.551 | 0.460 | 0.503 | 0.537 |
| Smiles-Bert | 0.556 | 0.513 | 0.498 | 0.443 | 0.468 | 0.491 |
| TEINet | 0.520 | 0.370 | 0.980 | 0.350 | 0.510 | 0.470 |
| ChemBERTa | 0.649 | 0.614 | 0.542 | 0.446 | 0.488 | 0.511 |
| TITAN | 0.525 | 0.660 | 0.537 | 0.516 | 0.523 | 0.551 |
| ERGO II | 0.512 | 0.643 | 0.503 | 0.516 | 0.534 | 0.503 |
| NetTCR | 0.534 | 0.524 | 0.557 | 0.563 | 0.609 | 0.524 |
| DlpTcr | 0.564 | 0.688 | 0.951 | 0.692 | 0.801 | 0.838 |
| Imrex | 0.660 | 0.685 | 0.113 | 0.731 | 0.194 | 0.518 |
| **Original NA** |  |  |  |  |  |  |
| TIDE | 0.901 | 0.964 | 0.997 | 0.966 | 0.981 | 0.995 |
| Smiles-Bert | 0.658 | 0.705 | 0.637 | 0.890 | 0.821 | 0.928 |
| TEINet | 0.972 | 0.951 | 0.991 | 0.963 | 0.981 | 1.000 |
| ChemBERTa | 0.731 | 0.972 | 0.994 | 0.977 | 0.986 | 0.993 |
| TITAN | 0.820 | 0.980 | 0.999 | 0.990 | 0.990 | 1.000 |
| ERGO II | 0.623 | 0.689 | 0.621 | 0.857 | 0.603 | 0.725 |
| NetTCR | 0.593 | 0.602 | 0.615 | 0.813 | 0.498 | 0.632 |
| DlpTcr | 0.666 | 0.977 | 0.336 | 0.553 | 0.389 | 0.474 |
| Imrex | 0.760 | 0.954 | 0.987 | 0.965 | 0.976 | 0.988 |
| **Original RN** |  |  |  |  |  |  |
| TIDE | 0.570 | 0.620 | 0.260 | 0.400 | 0.320 | 0.400 |
| Smiles-Bert | 0.573 | 0.602 | 0.521 | 0.763 | 0.702 | 0.817 |
| TEINet | 0.531 | 0.417 | 0.880 | 0.351 | 0.504 | 0.353 |
| ChemBERTa | 0.530 | 0.601 | 0.213 | 0.339 | 0.226 | 0.352 |
| TITAN | 0.501 | 0.660 | 0.413 | 0.562 | 0.431 | 0.347 |
| ERGO II | 0.521 | 0.505 | 0.537 | 0.689 | 0.531 | 0.625 |
| NetTCR | 0.535 | 0.511 | 0.547 | 0.621 | 0.509 | 0.662 |
| DlpTcr | 0.501 | 0.658 | 0.991 | 0.660 | 0.792 | 0.829 |
| Imrex | 0.562 | 0.657 | 0.026 | 0.387 | 0.046 | 0.377 |

A comparison between the **Generated** and **Original** datasets—namely *Gen NA* vs. *Original NA* and *Gen RN* vs. *Original RN*—reveals that models trained on generated datasets generally achieve better performance. For example, LANTERN attains an AUC-ROC of 0.88 on *Gen NA* compared to 0.90 on *Original NA*, and 0.66 on *Gen RN* compared to 0.57 on *Original RN*. This trend suggests that the negative sample generation strategy enhances training by providing a more balanced and challenging learning environment that better reflects real-world complexities.

By generating diverse and realistic negative samples in the *Gen NA* and *Gen RN* datasets, the models are exposed to a broader spectrum of binding and non-binding interactions. This exposure strengthens the models’ ability to capture subtle distinctions in TCR–peptide binding patterns, thereby improving both prediction accuracy and robustness.

#### Challenging Nature of the Random Control Setting

Although TIDE achieves an AUC-ROC of 0.66 under the random control setting, this scenario remains highly challenging. Random control generates negative samples rapidly and cost-effectively, but the inherent stochasticity produces noisier training signals than the reference control setting. Some randomly sampled non-binding pairs may partially resemble true binders, increasing the ambiguity and difficulty of distinguishing positives from negatives. While the random control setting is attractive for its scalability, its variability underscores the need for more robust strategies to improve model reliability.

### D. Attention-Weighted Embedding Similarity Visualization

To assess whether the attention mechanism enhances the discriminative power of TCR–peptide embeddings, we visualize the pairwise similarity distribution of cross-attention-weighted representations.

Figure 4 illustrates a heatmap of attention-weighted cosine similarity between TCR and peptide embeddings obtained from the cross-attention module. To construct this visualization, the 50 most frequent peptides and 50 most frequent TCRs from the Gen NA test set are selected to form a 50 *×* 50 similarity matrix. Each similarity score is normalized to the [0, 1] range using min–max scaling, and ground-truth positive pairs are highlighted with red markers.

**Fig. 4.**
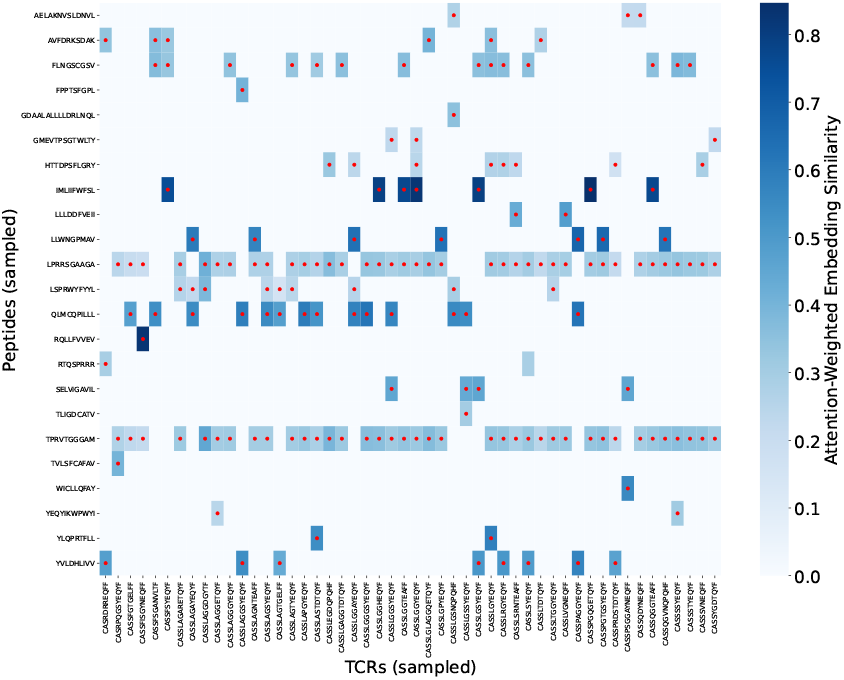
Heatmap of cross-attention-weighted embedding similarity (scaled to [0, 1]) between TCRs and peptides. Red markers indicate positive (binding) pairs concentrated in high-similarity regions.

The heatmap reveals that positive TCR–peptide pairs are largely concentrated in high-similarity regions. This pattern indicates that the attention mechanism successfully captures meaningful interaction patterns, creating a more separable embedding space that facilitates accurate classification.

### E. Embedding Clustering of the Pretraining Model

To evaluate the quality and clustering tendency of the learned TCR–peptide embeddings, we adopt four widely used clustering metrics [42]: Hopkins Score (HS), Cluster Tendency Score (CTS), Silhouette Index (SI), and Calinski–Harabasz Index (CHI). HS and CTS measure how likely the embeddings form meaningful clusters rather than a uniform distribution:

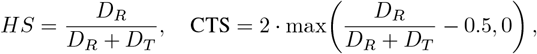

where *D*_*T*_ and *D*_*R*_ are the nearest-neighbor distances of sampled and random points, respectively. SI evaluates the cohesion and separation of clusters as

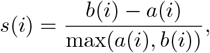

where *a*(*i*) and *b*(*i*) are the intra- and nearest-intercluster distances for point *i*. Finally, CHI measures the ratio of between- to within-cluster dispersion,

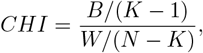

where *B* and *W* denote the between- and within-cluster variances, *K* is the number of clusters, and *N* is the total number of points. Together, these metrics capture both clustering tendency and structural quality of the learned embeddings.

The results are shown in Figure 5. Embeddings produced by the pretrained model exhibit markedly stronger clustering behavior than those without pretraining. The CTS reaches 0.95 with pretraining, compared to 0.41 without, reflecting a clear increase in the tendency to form clusters. Similarly, the HS rises from 0.61 to 0.84, indicating that the pretrained embeddings form more coherent clusters. The SI improves from 0.50 to 0.57, demonstrating that clusters become more well-defined, while the CHI increases from a scaled value of 0.9 (originally 2.12) to 1.0 (originally 389.06), showing substantially tighter and better-separated clusters after pretraining.

**Fig. 5.**
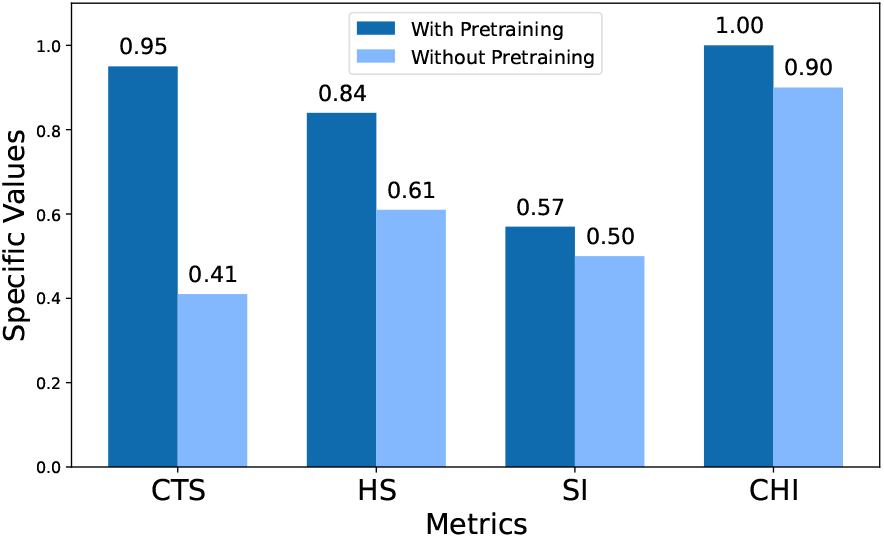
Comparison of clustering metrics for embeddings with and without pretraining. SI and CHI scores are scaled with a sigmoid function for readability.

These results collectively highlight the benefits of incorporating pretraining in TIDE. The significant gains across CTS, HS, SI, and CHI indicate that pretraining produces higher-quality embeddings with clearer cluster structures, which enhances the model’s ability to capture discriminative TCR–peptide interaction patterns and ultimately improves prediction robustness.

### F. Few-Shot Learning Capabilities of TIDE

Building on the strong zero-shot performance of TIDE, its ability to operate in few-shot settings is further examined. Fewshot learning is particularly relevant for practical applications where experimental TCR–peptide binding data is scarce. In this setting, the model is trained on only a handful of samples per peptide, allowing us to assess its adaptability and generalization with minimal supervision. To evaluate this capability, we control the number of TCR–peptide pairs available for each peptide.

Figure 6 presents results where 20% of the peptides in the test set are included in the training set, and the number of TCR–peptide pairs per peptide is gradually increased. The results show that TIDE rapidly improves with just a few examples per peptide (fewer than five), demonstrating its ability to learn binding patterns from extremely limited data. As the number of pairs grows, performance gains taper off, suggesting that the model reaches a saturation point where additional training data yields diminishing returns.

**Fig. 6.**
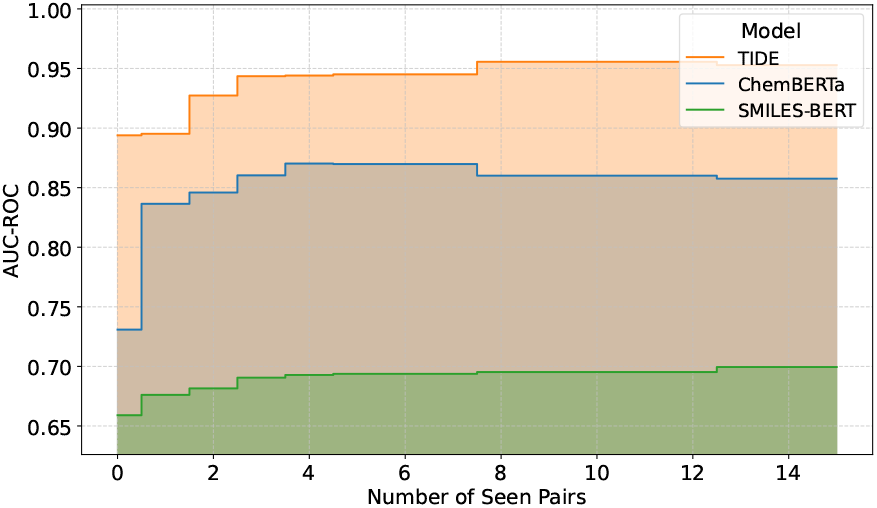
Few-shot learning performance of TIDE.

## V. Discussion and Conclusion

TIDE advances TCR–pMHC binding prediction by integrating protein and molecular language models with a cross-attention fusion mechanism. Leveraging ESM and MolFormer embeddings allows the model to capture both sequence-level and chemical representations, achieving strong performance in zero-shot and few-shot settings, particularly for previously unseen epitopes. A central strength of TIDE lies in its ability to generalize from limited data, consistently outperforming both traditional deep learning models and recent LLM-based methods. The multi-head cross-attention module enhances interpretability by aligning TCR and peptide features while improving discriminative capability.

Overall, TIDE represents a step forward in TCR–pMHC interaction prediction. Its capacity to generalize across novel epitopes and maintain strong performance under low-data conditions positions it as a valuable tool for supporting the development of personalized immunotherapies and vaccines. Future work will focus on incorporating richer biological context to further improve accuracy and interpretability.

## References

[1] M. M. Davis and P. J. Bjorkman, “T-cell antigen receptor genes and t-cell recognition,” Nature, vol. 334, no. 6181, pp. 395–402, 1988.

[2] M. Krogsgaard and M. M. Davis, “How t cells’ see’antigen,” Nature immunology, vol. 6, no. 3, pp. 239–245, 2005.

[3] R.-Y. Chen, Y. Zhu, Y.-Y. Shen, Q.-Y. Xu, H.-Y. Tang, N.-X. Cui, L. Jiang, X.-M. Dai, W.-Q. Chen, Q. Lin et al., “The role of pd-1 signaling in health and immune-related diseases,” Frontiers in Immunology, vol. 14, p. 1163633, 2023.

[4] Q. Tang, Y. Chen, X. Li, S. Long, Y. Shi, Y. Yu, W. Wu, L. Han, and S. Wang, “The role of pd-1/pd-l1 and application of immune-checkpoint inhibitors in human cancers,” Frontiers in immunology, vol. 13, p. 964442, 2022.

[5] M. Cai, S. Bang, P. Zhang, and H. Lee, “Atm-tcr: Tcr-epitope binding affinity prediction using a multi-head self-attention model,” Frontiers in immunology, vol. 13, p. 893247, 2022.

[6] L. Heumos, A. C. Schaar, C. Lance, A. Litinetskaya, F. Drost, L. Zappia, M. D. Lücken, D. C. Strobl, J. Henao, F. Curion et al., “Best practices for single-cell analysis across modalities,” Nature Reviews Genetics, vol. 24, no. 8, pp. 550–572, 2023.

[7] I. Springer, N. Tickotsky, and Y. Louzoun, “Contribution of t cell receptor alpha and beta cdr3, mhc typing, v and j genes to peptide binding prediction,” Frontiers in immunology, vol. 12, p. 664514, 2021.

[8] C. Qi, H. Fang, T. Hu, W. Zhi et al., “Enhancing tcr-peptide interaction prediction with pretrained language models and molecular representations,” arXiv preprint arXiv:2505.01433, 2025.

[9] C. Qi, Y. Chen, J. Zhang, and W. Zhi, “Clustering with communication: A variational framework for single cell representation learning,” arXiv preprint arXiv:2505.04891, 2025.

[10] C. Qi, H. Fang, T. Hu, S. Jiang, and W. Zhi, “Bidirectional mamba for single-cell data: Efficient context learning with biological fidelity,” arXiv preprint arXiv:2504.16956, 2025.

[11] L. V. Castorina, F. Grazioli, P. Machart, A. Mösch, and F. Errica, “Assessing the generalization capabilities of tcr binding predictors via peptide distance analysis,” bioRxiv, pp. 2023–07, 2023.

[12] T. Mayassi, L. B. Barreiro, J. Rossjohn, and B. Jabri, “A multilayered immune system through the lens of unconventional t cells,” Nature, vol. 595, no. 7868, pp. 501–510, 2021.

[13] J. A. Pai and A. T. Satpathy, “High-throughput and single-cell t cell receptor sequencing technologies,” Nature methods, vol. 18, no. 8, pp. 881–892, 2021.

[14] J. Lipkova, R. J. Chen, B. Chen, M. Y. Lu, M. Barbieri, D. Shao, A. J. Vaidya, C. Chen, L. Zhuang, D. F. Williamson et al., “Artificial intelligence for multimodal data integration in oncology,” Cancer cell, vol. 40, no. 10, pp. 1095–1110, 2022.

[15] M. Fidelle, C. Rauber, C. Alves Costa Silva, A.-L. Tian, I. Lahmar, A.-L. M. de La Varende, L. Zhao, C. Thelemaque, I. Lebhar, M. Messaoudene et al., “A microbiota-modulated checkpoint directs immunosuppressive intestinal t cells into cancers,” Science, vol. 380, no. 6649, p. eabo2296, 2023.

[16] J.-W. Sidhom, H. B. Larman, D. M. Pardoll, and A. S. Baras, “Deeptcr is a deep learning framework for revealing sequence concepts within t-cell repertoires,” Nature communications, vol. 12, no. 1, p. 1605, 2021.

[17] A. Weber, J. Born, and M. Rodriguez Martínez, “Titan: T-cell receptor specificity prediction with bimodal attention networks,” Bioinformatics, vol. 37, no. Supplement 1, pp. i237–i244, 2021.

[18] T. Lu, Z. Zhang, J. Zhu, Y. Wang, P. Jiang, X. Xiao, C. Bernatchez, J. V. Heymach, D. L. Gibbons, J. Wang et al., “Deep learning-based prediction of the t cell receptor–antigen binding specificity,” Nature machine intelligence, vol. 3, no. 10, pp. 864–875, 2021.

[19] F. Grazioli, A. Mösch, P. Machart, K. Li, I. Alqassem, T. J. O’Donnell, and M. R. Min, “On tcr binding predictors failing to generalize to unseen peptides,” Frontiers in immunology, vol. 13, p. 1014256, 2022.

[20] Y. Jiang, M. Huo, and S. Cheng Li, “Teinet: a deep learning framework for prediction of tcr–epitope binding specificity,” Briefings in bioinformatics, vol. 24, no. 2, p. bbad086, 2023.

[21] D. Korpela, E. Jokinen, A. Dumitrescu, J. Huuhtanen, S. Mustjoki, and H. Lähdesmäki, “Epic-trace: predicting tcr binding to unseen epitopes using attention and contextualized embeddings,” Bioinformatics, vol. 39, no. 12, p. btad743, 2023.

[22] K. E. Wu, K. Yost, B. Daniel, J. Belk, Y. Xia, T. Egawa, A. Satpathy, H. Chang, and J. Zou, “Tcr-bert: learning the grammar of t-cell receptors for flexible antigen-binding analyses,” in Machine Learning in Computational Biology. PMLR, 2024, pp. 194–229.

[23] V. Mhanna, H. Bashour, K. Lê Quý, P. Barennes, P. Rawat, V. Greiff, and E. Mariotti-Ferrandiz, “Adaptive immune receptor repertoire analysis,” Nature Reviews Methods Primers, vol. 4, no. 1, p. 6, 2024.

[24] Y. Xu, X. Liu, X. Cao, C. Huang, E. Liu, S. Qian, X. Liu, Y. Wu, F. Dong, C.-W. Qiu et al., “Artificial intelligence: A powerful paradigm for scientific research,” The Innovation, vol. 2, no. 4, 2021.

[25] T. Fan, M. Zhang, J. Yang, Z. Zhu, W. Cao, and C. Dong, “Therapeutic cancer vaccines: advancements, challenges and prospects,” Signal Transduction and Targeted Therapy, vol. 8, no. 1, p. 450, 2023.

[26] A. Rives, J. Meier, T. Sercu, S. Goyal, Z. Lin, J. Liu, D. Guo, M. Ott, C. L. Zitnick, J. Ma et al., “Biological structure and function emerge from scaling unsupervised learning to 250 million protein sequences,” Proceedings of the National Academy of Sciences, vol. 118, no. 15, p. e2016239118, 2021, bioRxiv 10.1101/622803. [Online]. Available: https://www.pnas.org/doi/full/10.1073/pnas.2016239118

[27] D. Weininger, “Smiles, a chemical language and information system. 1. introduction to methodology and encoding rules,” Journal of chemical information and computer sciences, vol. 28, no. 1, pp. 31–36, 1988.

[28] J. Ross, B. Belgodere, V. Chenthamarakshan, I. Padhi, Y. Mroueh, and P. Das, “Large-scale chemical language representations capture molecular structure and properties,” Nature Machine Intelligence, vol. 4, no. 12, pp. 1256–1264, 2022.

[29] Z. Lin, H. Akin, R. Rao, B. Hie, Z. Zhu, W. Lu, A. dos Santos Costa, M. Fazel-Zarandi, T. Sercu, S. Candido et al., “Language models of protein sequences at the scale of evolution enable accurate structure prediction,” BioRxiv, vol. 2022, p. 500902, 2022.

[30] N. L. La Gruta, S. Gras, S. R. Daley, P. G. Thomas, and J. Rossjohn, “Understanding the drivers of mhc restriction of t cell receptors,” Nature Reviews Immunology, vol. 18, no. 7, pp. 467–478, 2018.

[31] V. I. Jurtz, L. E. Jessen, A. K. Bentzen, M. C. Jespersen, S. Mahajan, R. Vita, K. K. Jensen, P. Marcatili, S. R. Hadrup, B. Peters et al., “Nettcr: sequence-based prediction of tcr binding to peptide-mhc complexes using convolutional neural networks,” BioRxiv, p. 433706, 2018.

[32] A. Montemurro, V. Schuster, H. R. Povlsen, A. K. Bentzen, V. Jurtz, W. D. Chronister, A. Crinklaw, S. R. Hadrup, O. Winther, B. Peters et al., “Nettcr-2.0 enables accurate prediction of tcr-peptide binding by using paired tcrα and β sequence data,” Communications biology, vol. 4, no. 1, p. 1060, 2021.

[33] K. Singhal, S. Azizi, T. Tu, S. S. Mahdavi, J. Wei, H. W. Chung, N. Scales, A. Tanwani, H. Cole-Lewis, S. Pfohl et al., “Large language models encode clinical knowledge,” Nature, vol. 620, no. 7972, pp. 172– 180, 2023.

[34] M. Jin, S. Wang, L. Ma, Z. Chu, J. Y. Zhang, X. Shi, P.-Y. Chen, Y. Liang, Y.-F. Li, S. Pan et al., “Time-llm: Time series forecasting by reprogramming large language models,” arXiv preprint arXiv:2310.01728, 2023.

[35] S. Wang, Y. Guo, Y. Wang, H. Sun, and J. Huang, “Smiles-bert: large scale unsupervised pre-training for molecular property prediction,” in Proceedings of the 10th ACM international conference on bioinformatics, computational biology and health informatics, 2019, pp. 429–436.

[36] S. Chithrananda, G. Grand, and B. Ramsundar, “Chemberta: large-scale self-supervised pretraining for molecular property prediction,” arXiv preprint arXiv:2010.09885, 2020.

[37] Z. Xu, M. Luo, W. Lin, G. Xue, P. Wang, X. Jin, C. Xu, W. Zhou, Y. Cai, W. Yang et al., “Dlptcr: an ensemble deep learning framework for predicting immunogenic peptide recognized by t cell receptor,” Briefings in Bioinformatics, vol. 22, no. 6, p. bbab335, 2021.

[38] P. Moris, J. De Pauw, A. Postovskaya, S. Gielis, N. De Neuter, W. Bittremieux, B. Ogunjimi, K. Laukens, and P. Meysman, “Current challenges for unseen-epitope tcr interaction prediction and a new perspective derived from image classification,” Briefings in bioinformatics, vol. 22, no. 4, p. bbaa318, 2021.

[39] D. Feng, C. J. Bond, L. K. Ely, J. Maynard, and K. C. Garcia, “Structural evidence for a germline-encoded t cell receptor–major histocompatibility complex interaction’codon’,” Nature immunology, vol. 8, no. 9, pp. 975– 983, 2007.

[40] J. Glanville, H. Huang, A. Nau, O. Hatton, L. E. Wagar, F. Rubelt, X. Ji, A. Han, S. M. Krams, C. Pettus et al., “Identifying specificity groups in the t cell receptor repertoire,” Nature, vol. 547, no. 7661, pp. 94–98, 2017.

[41] L. Rowen, B. F. Koop, and L. Hood, “The complete 685-kilobase dna sequence of the human β t cell receptor locus,” Science, vol. 272, no. 5269, pp. 1755–1762, 1996.

[42] G. Zheng, J. Rymuza, E. Gharavi, N. J. LeRoy, A. Zhang, and N. C. Sheffield, “Methods for evaluating unsupervised vector representations of genomic regions,” NAR Genomics and Bioinformatics, vol. 6, no. 3, p. lqae086, 2024.

